# High congruence of karyotypic and molecular data on *Hypostomus* species from the Paraná River basin

**DOI:** 10.1101/2020.09.22.308437

**Authors:** Dinaíza Abadia Rocha-Reis, Rubens Pasa, Karine Frehner Kavalco

## Abstract

The Hypostomini tribe comprises a single genus, *Hypostomus*, which possibly contains several monophyletic groups because of significant morphological variation and a variety of diploid numbers and karyotype formulas. The objective of this study was to infer evolutionary relationships among some species of *Hypostomus* found in the Paraná River basin and subsequently to identify chromosomal synapomorphies in the groupings formed. Two nuclear genes, *rag1* and *rag2*, and two mitochondrial genes, *mt-co1* and *mt-cyb*, were used to establish evolutionary relationships. Phylogenetic trees were inferred using the maximum likelihood (ML) method for *mt-co1* and Bayesian analysis (BA) for all genes concatenated. Both phylogenetic trees showed two large monophyletic clades within *Hypostomus*. These clades are based on chromosome number, where haplogroup I contains individuals with 66–68 chromosomes, and haplogroup II contains species with 72–80 chromosomes. A third monophyletic haplogroup was also observed using ML, formed by *H. faveolus* and *H. cochliodon*, which present 2n = 64, reinforcing the separation of groups in *Hypostomus* by diploid number. Robertsonian rearrangements were responsible for forming the different diploid numbers and for the diversity of karyotype formulas. The groups based on traditional morphological taxonomy are considered artificial in this study; the staining pattern, which separates the two large groups morphologically and is supported by little chromosomal evidence, was instead determined to show homoplasy. Ag-NORs are predominantly multiple and located on st/a chromosomes, along with 18S rDNA sites; 5S rDNA sites are often seen in an interstitial position, following the trend already described for vertebrates.

## 1. Introduction

The Hypostominae subfamily (Siluriformes, Loricariidae) comprises approximately 500 valid species (Eschmeyer and Fong 2019), and shows great variation in coloration and external morphology (Oyakawa et al. 2005; Zawadzki et al. 2008). Armbruster (2004) divided this subfamily into five tribes: Corymbophanini, Rhinelepini, Hypostomini, Ancistrini and Pterygoplichthyini. The Hypostomini tribe comprises only the genus *Hypostomus* Lacépède 1803 (Armbruster 2004), and has a large number of cytogenetic studies (Appendix A).

Based on its great species diversity (Muller and Weber 1992; Armbruster 2004; Zawadzki et al. 2004) and on the identification of a variety of diploid numbers and karyotype formulas (Appendix A), it is suggested that several monophyletic groups may be found within *Hypostomus*.

These fish, popularly known as plecos, cascudos (in their native range), or armored catfish have a wide geographic distribution, non-migratory behavior, and high adaptive performance, favoring the formation of populations even without physical barriers (Alves et al. 2005; Bickford et al. 2007). Thus, studies on *Hypostomus* populations integrating cytogenetic and molecular tools can provide valuable information on phylogeography and microevolutionary processes.

A similar approach was used for some species of the characid genus *Astyanax* Baird & Girard (1854) (Pazza et al. 2018). These authors showed that characteristics such as the quantity and position of chromosomal markers, as the 5S rDNA and the As-51 satellite DNA, were related to the divergence of large groups of species and species complexes, as well as macrostructural karyotype characteristics.

The objective of this study was, therefore, to outline the phylogenetic and evolutionary relationships of some *Hypostomus* species found in the Alto Paraná River basin to identify possible relationships between chromosomal and molecular variations in this group.

## 2. Material and methods

*Hypostomus* specimens were collected in the Grande, Paranaíba, Paranapanema, and Tietê river basins, totaling 65 individuals belonging to *H. ancistroides* (Ihering, 1911), *H. faveolus* (Zawadzki et al. 2008), *H. margaritifer* (Regan, 1908), *H. paulinus* (Ihering, 1905), *H. regani* (Ihering, 1905), *Hypostomus* sp., *Hypostomus* sp. 2, *H. strigaticeps* (Regan, 1908), and *H. tietensis* (Ihering, 1905) (Appendix B).

Samples of liver and heart tissues from the specimens collected were used to obtain genomic DNA. Appropriate kits were used for extractions, following the manufacturer’s recommendations (PureLink™ Genomic DNA Kit, Invitrogen, Thermo Fisher Scientific, Life Technologies, Carlsbad, CA, USA).

Sequences of two mitochondrial genes, the cytochrome oxidase subunit I (*mt-co1*) and cytochrome b (*mt-cyb*), and two nuclear genes, the recombination-activating gene 1 (*rag1*) and recombination-activating gene 2 (*rag2*), were amplified. The primers used for *mt-co1* were Fish F1 (5’TCAACCAACCACAAAGACATTGGC-3’) and Fish R1 (5’TAGACTTCTGGGTGGCCAAAGA-3’). Primers CytbFc and CytbRc were used for *mt-cyb*, RAG1Fa and RAG1R1186 for *rag1*, and RAG2Fc and RAG2R196 for *rag2*, as described by Lujan et al. (2015).

The PCR was conducted with cycles of 95°C for two min, followed by 35 cycles of 94°C for 30 s, X°C for 30 s (depending on primer pairs), and 72°C for 1 min, with a final extension of 72°C for 10 min. After the end of the reaction, the amplified fragments were checked by electrophoresis in a 1% agarose gel. If positive, they were sent for sequencing by a third-party company.

The sequences obtained were edited using the CodonCode Aligner 6.0.2 software, verified in GenBank (http://www.ncbi.nlm.nih.gov) using the BLASTN tool, and aligned using the ClustalW 1.6 algorithm (Thompson et al. 1994) implemented in MEGA 7.0.21 software (Kumar et al. 2016).

To reconstruct phylogenetic trees, *mt-co1* sequences from other species of *Hypostomus* were added (all available from GenBank), and *Pterygoplichthys* species were used as outgroups (Appendix C).

Maximum likelihood (ML) analyses were conducted separately for each gene with IQ-TREE 1.5.6 software (Nguyen et al. 2015; Chernomor et al. 2016). The best nucleotide substitution model was estimated by the Bayesian information criterion (BIC). The analyses were carried out with 1,000 bootstrap replicates.

Bayesian analysis (BA) was conducted using MrBayes 3.2.6 software (Ronquist et al. 2012) with gene concatenation. An independent search for the best nucleotide substitution model was carried out for each gene using PartitionFinder 1.1.1 software (Lanfear et al. 2012). After running 50 million Markov Chain Monte Carlo simulations, the length of sampling chain per thousand generations was evaluated using Tracer 1.7 software (Rambaut et al. 2018) to verify the effective sample size and chain convergence. The Tree Annotator 1.8 software discarded the first 25% of trees as burn-in.

The visualization of trees (ML and BA) was carried out using FigTree 1.4.2 software (Rambaut et al. 2012).

## 3. Results

The ML trees obtained by the genes *mt-cyb, rag1*, and *rag2* presented many polytomies and an absence of group formation (data not shown) and were therefore not informative for the phylogeny of the genus *Hypostomus*.

However, the ML tree obtained using *mt-co1* showed the formation of three groups within *Hypostomus* (Fig. 1 and Fig. 2). This pattern was similar to that obtained in the Bayesian tree with sequence concatenation (Fig. 3 and Fig. 4). The first lineage (Haplogroup I – in green) comprised *H. ancistroides, H*. cf. *tietensis, H. affinis, H. commersoni*, and *H. derbyi*. The second lineage (Haplogroup II – in orange) comprised *H. regani, H. strigaticeps, H. paulinus, H. margaritifer, Hypostomus* sp., *H. hermanni*, *H. iheringii*, *H. nigromaculatus*, and *H. topavae*. Haplogroup III (Fig. 1 – in blue) was presented as monophyletic in ML, but not in BA (Fig. 3), and comprised only *H. faveolus* and *H. cochliodon*. *Hypostomus plecostomus* did not group with other species of *Hypostomus*; it was related to species of an external group from the genus *Pterigoplichthys* (tribe Pterygoplichthyini).

**Fig. 1.**
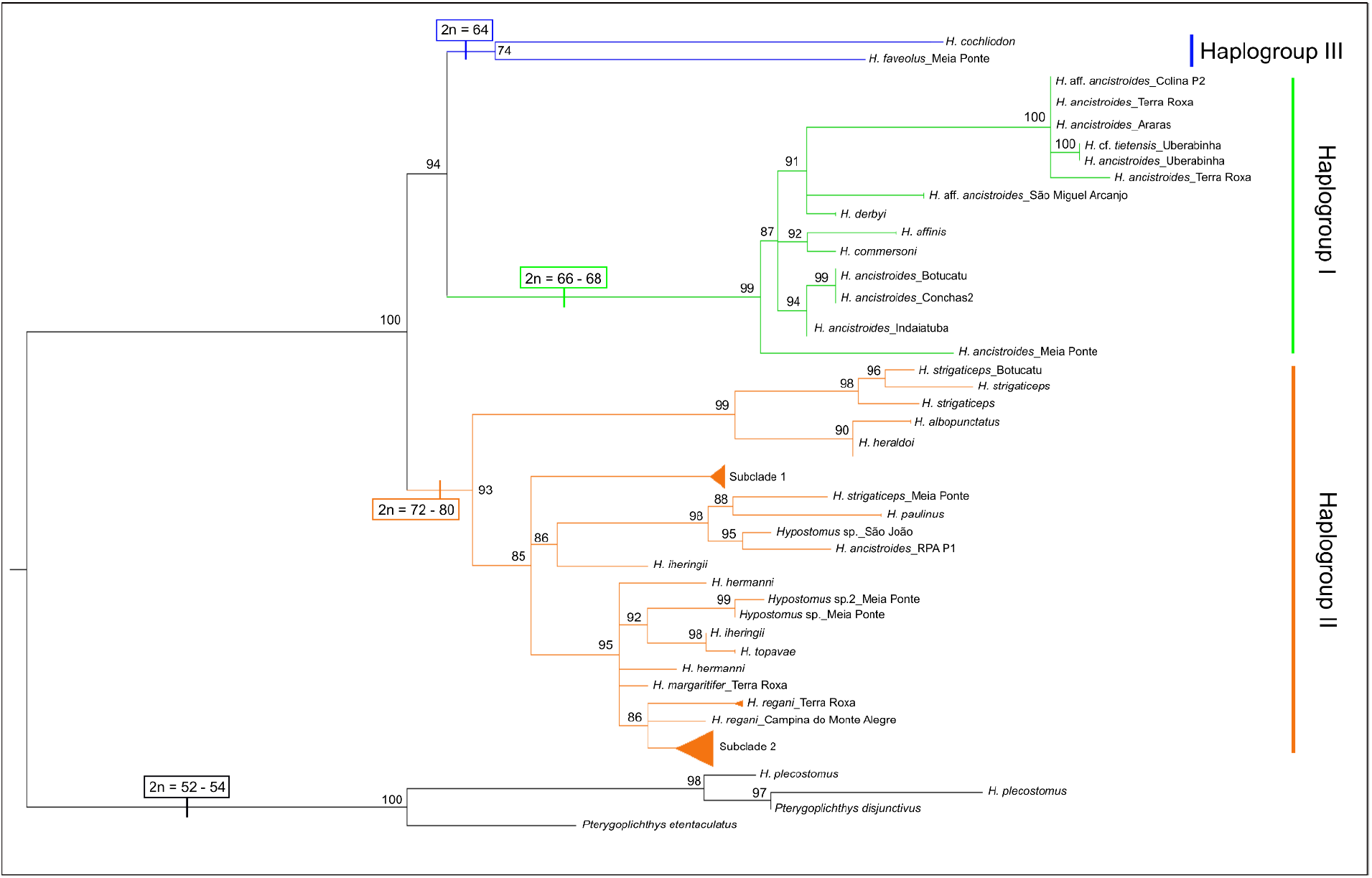
Maximum Likelihood tree obtained for the species of *Hypostomus* through *mt-co1* sequences. The numbers on the nodes represent the bootstrap values

**Fig. 2.**
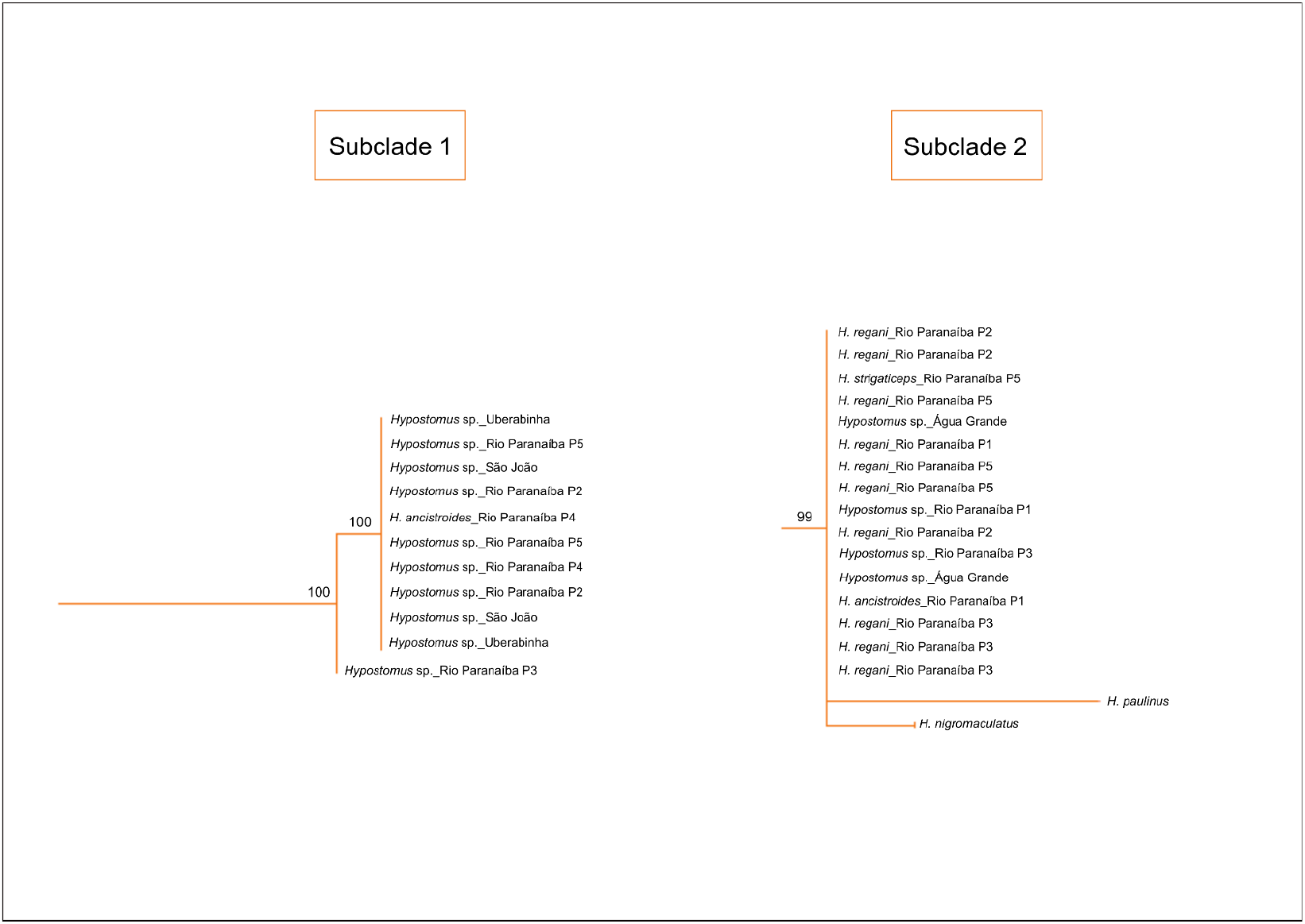
Details of the Operational Taxonomic Units (OTUs) present in Subclades 1 and 2 in the Maximum Likelihood Analysis with *mt-co1* sequences. The numbers on the nodes represent the bootstrap values

**Fig. 3.**
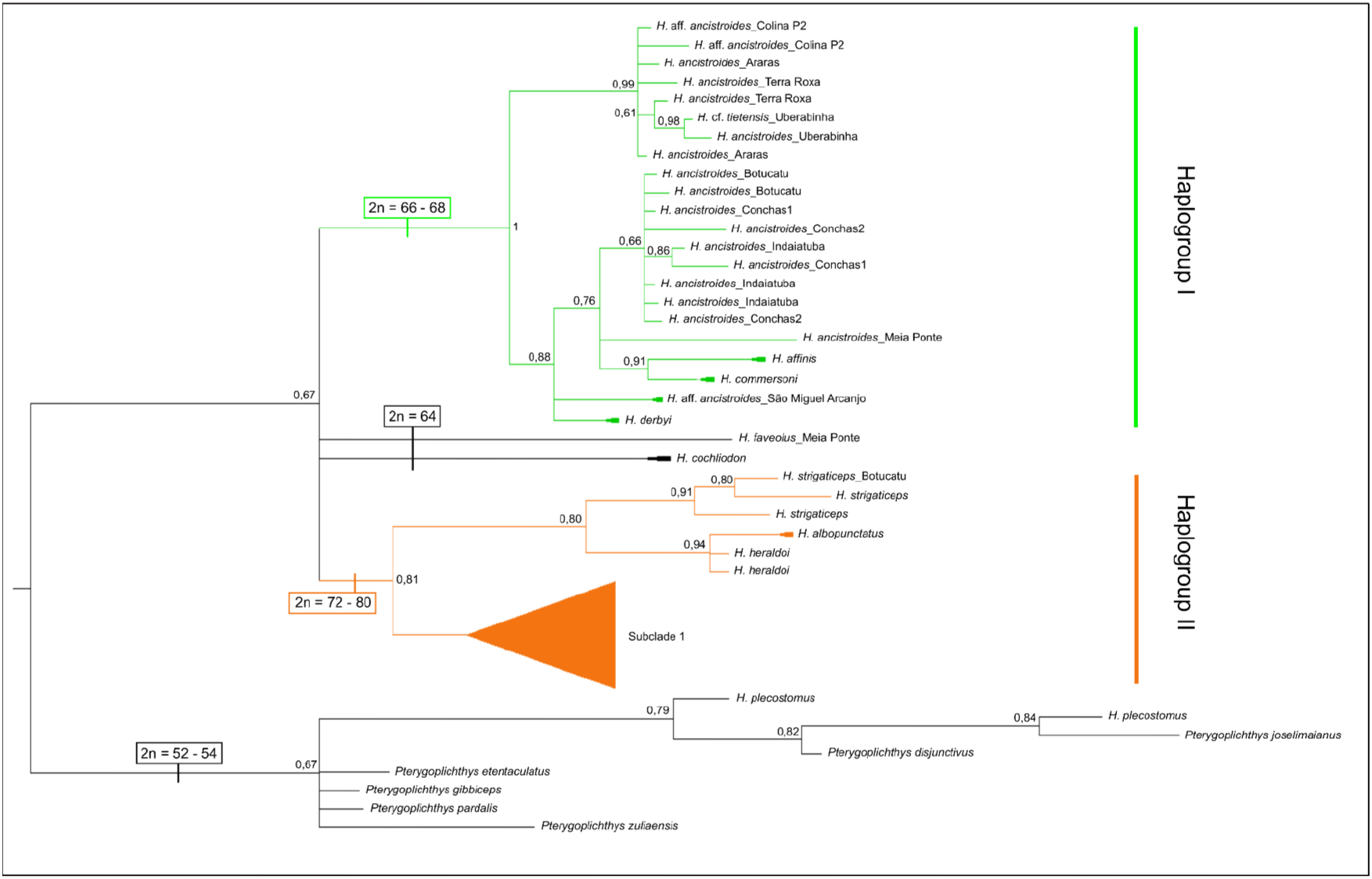
Bayesian tree obtained for *Hypostomus* species from the concatenation of nuclear and mitochondrial sequences. The numbers on the nodes represent the posterior probability values

**Fig. 4.**
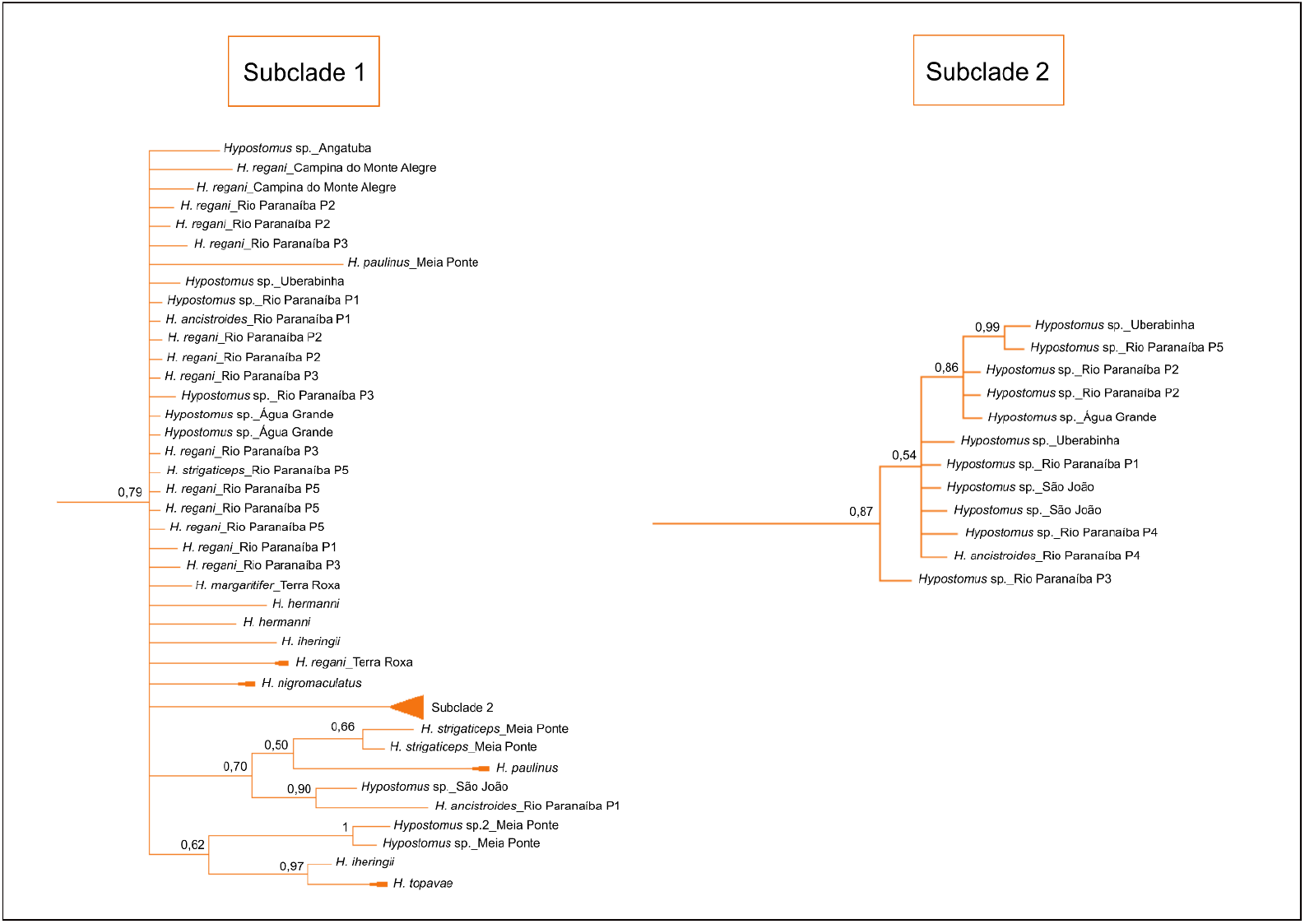
Details of the Operational Taxonomic Units (OTUs) present in Subclades 1 and 2 in the Bayesian Analysis with concatenation of nuclear and mitochondrial sequences. The numbers on the nodes represent the posterior probability values

## 4. Discussion

The present results showed the formation of at least two large monophyletic groups in *Hypostomus* (Fig. 1 and Fig. 3), related to the diploid number of the species. In both analysis, Haplogroup I contained species with 66–68 chromosomes, whereas Haplogroup II contained species with 72–80 chromosomes (already observed by Alves et al. 2006; Bueno et al. 2014). The phylogenetic analyses with ML also showed the formation of Haplogroup III – monophyletic (Fig. 1, in blue), containing species with 2n = 64 chromosomes, reinforcing the separation of groups in *Hypostomus* by diploid number.

Despite the formation of large groups, the trees still have many unresolved relationships within these clades. Montoya-Burgos et al. (1997, 1998, 2002) obtained similar results using 12S and 16S mtDNA rRNA and D-loop, with monophyletic clades well established within *Hypostomus*, but the relationship among species within the clades was unresolved and poorly supported. Thus, there is a need to sample more taxa of the genus *Hypostomus* to infer a more robust phylogeny.

The ML trees obtained from *mt-cyb*, *rag1*, and *rag2* further evidenced these unresolved relationships within the clades; thus, they were considered not informative for the genus. In addition to the matter of representativeness, this may have occurred because of the mutation rates of the genes used. The *rag1* and *rag2* trees presented the highest number of polytomies because nuclear genes have a low evolution rate compared to mitochondrial genes (Avise *et al*., 1987). Despite being mitochondrial, *mt-cyb* is considered conserved and is used most often in taxa studies above the species level (Pereira 2000). Therefore, *mt-co1* was selected as the best gene for recovering phylogenetic relationships among species. In future studies, approaches using the mtDNA control region (D-loop) may rescue the most recent evolutionary history of the group.

Muller and Weber (1992) suggested the existence of two groups in *Hypostomus* based on the morphology and body coloration: (1) the *plecostomus* group, which have dark spots on a light-colored body, medium-sized jaw, and small crowned teeth; and (2) the *regani* group, which have clear spots on a dark-colored body, large jaws, and large crowned teeth. Zawadzki et al. (2004) and Alves et al. (2006) observed the corresponding groups with cytogenetic data: the *plecostomus* group had 66–68 chromosomes, and the *regani* group had 72–74 chromosomes.

In our analyses, however, the groupings were less clear, since there are species of the *plecostomus* group belonging to the clade with species that present more than 72 chromosomes, such as *H. nigromaculatus*, *H. topavae*, *H. iheringii*, *H. hermanni*, and *H. paulinus*, showing that the karyotype data may be correlated with the division of monophyletic groups in *Hypostomus*, and that the use of morphology to separate species shows the formation of polyphyletic or artificial groups. In this way, the staining pattern, which separates the two large groups morphologically and is supported by little chromosomal evidence, shows homoplasy.

Two inconsistencies were found in both phylogenetic trees in Haplogroup II Subclades 1 and 2 (Figs.1–4). There are two individuals of *H. ancistroides* from Rio Paranaíba P1 and Rio Paranaíba P2 (Fig. 2 and Fig. 4). It is believed that this may be an identification mistake, as the lineage that gives rise to *H. ancistroides* belongs to group I (Fig. 1 and Fig. 3). Unfortunately, we could not confirm the identity of this individual by chromosome number due to the lack of cytogenetic material of the specimen.

The similar morphology among species may be the result of selective environmental pressure. *Hypostomus* species live on the bottom of rivers in sand banks and among rocks, and during drought, they shelter among rocks or submerged tree trunks (Weber 2003). The muddy habitat, combined with a scavenger’s diet based on algae and debris, could apply selective pressure to the body shape and color, being adaptive to the environment where they live. Thus, similar morphotypes are observed at different sites and in different lineages.

Although one of the morphological groups is named *plecostomus*, *H. plecostomus* was not grouped with the rest of the species from *Hypostomus* (Figs. 1 and Fig. 3). As the basis of group separation is focused on chromosome number, it was expected that *H. plecostomus* would be separated from the rest, since the original description of this species shows 54 chromosomes (Muramoto et al. 1968), the smallest diploid number of any species in the genus (Bueno et al. 2014). However, this species grouped with the outgroup, with a diploid number considered plesiomorphic. There are descriptions of *H*. prope *plecostomus* and *H*. cf. *plecostomus* with 68 chromosomes (Alves et al. 2012; Oliveira et al. 2015, respectively), showing that the group may include different taxa in addition to the one sampled in the present analysis.

In Loricariidae, 54 chromosomes is considered the basal condition (Artoni and Bertollo 2001). This has also been reported in species of Hypoptopomatinae (Andreata et al. 1993, 1994) and Loricariinae (Scavone and Julio Jr. 1995), as well as in the external group Trichomycteridae (Lima and Galetti Jr. 1990). Hypostominae, Rhinelepini (Artoni and Bertollo 2001), and Corymbophanini (Alves et al. 2005) present species with 54 chromosomes; Pterygoplichthyini and Ancistrini, considered sister groups (Armbruster 2004), present 2n = 52 chromosomes (Artoni and Bertollo 1996; 2001).

The diploid number of *Hypostomus* is much larger than those of other tribes of the same subfamily. The emergence of monophyletic clades based on chromosome number separation may be explained by large diploid number change events in ancestors. An event probably occurred resulting in an increase of 52–54 chromosomes (present in ancestors and sister groups) to 64–68 chromosomes, which later created the Haplogroups I and III (Fig. 1 and Fig. 3). Another chromosomal rearrangement event must have created the lineage of Haplogroup II (Fig. 1 and Fig. 3), changing the ancestors’ diploid number (64–68) to 72–84 chromosomes.

Artoni and Bertollo (1996) suggest that Robertsonian rearrangements, such as centric fissions and pericentric inversions, play a key role in Hypostominae evolution. According to these authors, there is an inverse relationship between the increase in diploid number and the proportion of chromosomes with two arms. This trend is confirmed in the species analyzed in this study, where there was an increase in subtelocentric/acrocentric chromosomes (st/a) with an increased diploid number (Appendix A), which can be clearly observed when comparing karyotype formulas among species with great diploid variation. A species of *Hypostomus* with 64 chromosomes, such as *H. faveolus*, presents 18m + 8sm + 22st + 16a (Bueno et al. 2013). With the increased diploid number, all chromosome types are also expected to increase. However, when we analyze the other extreme, *Hypostomus* sp. 2 with 84 chromosomes (6m + 16sm + 62st/a; Cereali et al. 2008), we see less metacentric chromosomes (m) and a significant increase in the number of st/a chromosomes, evidencing the existing fissions in this group.

Similar patterns are seen in both monophyletic groups when comparing the available cytogenetic data for *Hypostomus* species. The C-banding is distributed throughout the chromosomal complement of the species and is seen in all types of chromosomes (Appendix A). Although Artoni and Bertollo (1999) proposed the trend of heterochromatic blocks located in a pericentromeric region in *Hypostomus* species that have a greater diploid number, this cannot be seen in the available data. Either with a smaller or larger diploid number, the blocks are also found in the terminal region of long or short arms (Appendix A). This marker has not yet been extensively explored in population-based approaches for catfish, owing to the lack of apparent patterns among the data already available or the difficulty in obtaining satisfactory results for comparison, since most catfish chromosomes are type st/a and very small.

The phenotype of nucleolar organizer regions (NORs) shows simple and terminal sites that are plesiomorphic in Loricariidae (Artoni and Bertollo 1996; Oliveira and Gosztonyi 2000). Regarding the position of NORs, terminal NORs are conserved in *Hypostomus* and it is seen in all populations already described, varying only in which arm they are located (Appendix A). Regarding the number of sites, although simple NORs are observed in *Hypostomus* (Mendes-Neto et al. 2011; Endo et al. 2012; Bueno et al. 2013; Pansonato-Alves et al. 2013, among others), multiple NORs are the predominant phenotype in the genus (Appendix A). Another trend seen is the frequent location of these sites on st/a chromosomes that when multiple, have at least one pair of NORs located on this type of chromosome (Appendix A). In many populations, the Ag-NOR sites correspond to the C-banding. The presence of intercalated or adjacent heterochromatin segments with ribosomal sites is frequent and has already been reported by many authors (Kavalco et al. 2004; Rubert et al. 2008; Traldi et al. 2013). This association is believed to contribute to group evolution, since it allows the dispersion of NOR sites throughout the genome (Moreira-Filho et al. 1984; Vicari et al. 2008). Thus, polymorphisms involving heterochromatin may correspond to polymorphisms involving rDNAs.

Studies with GC- and AT-specific fluorochromes are not very common and account for about 28% and 20%, respectively, of all studies (Appendix A). The available data show that GC-specific fluorochromes are more common in *Hypostomus*; they are often coincident with NORs, showing that ribosomal sites may be interspersed with GC-rich regions. Data for AT-rich sites are scarce and often negative (Appendix A).

Data from 18S rDNA are still scarce when compared to data from classical cytogenetics, covering only 47.9% of descriptions. They coincide with the NOR markings in most cases (Appendix A), showing that most sites were active in the previous interphase. Thus, multiple sites are also predominant on st/a chromosomes in the terminal region, varying according to the arm carrying the sites.

The data for 5S rDNA in neotropical fish are even more scarce, as mentioned by Kavalco et al. (2004), and only 33.6% of the populations studied have descriptions for this marker (Appendix A). There is variation in site number: when they are in a simple phenotype, the predominant carrier chromosomes are m/sm; and when there are multiple sites, at least two of the sites (one pair) are also located in these two chromosomal types.

Regarding position, most of the 5S rDNA sites are in the pericentromeric or interstitial region. This pattern was previously reported by Martins and Galleti Jr. (2000) in Acipenseriformes, Anguilliformes, Characiformes, Perciformes, Salmoniformes, and Tetraodontiformes species, showing that the location of these sites may not be accidental, since this pattern is also seen in mammals and amphibians. Besides Hypostomini, in Hypostominae this phenotype was also observed in the Ancistrini (Mariotto et al. 2011; Silva 2014; Favarato et al. 2016), Rhinelepini (Silva 2014), and Pterygoplichthyini (Silva 2014) tribes. An interstitial distribution for 5S rDNA may then represent an advantage related to the organization of these genes in the vertebrate genome (Martins and Galleti Jr. 2000). A position inside the chromosomes could represent a form of “protection” for the gene if any type of rearrangement occurs, allowing more chances of maintenance and permanence of the gene transcriptional activity if there are changes in chromosome type. Such maintenance in a “protected” position also seems to be favorable in *Hypostomus*. Although few data are available for this marker, it is possible to delineate the trend of at least one chromosomal pair carrying either simple or multiple sites in a non-terminal position.

## 5. Conclusions

Phylogenetic analyses evidenced the formation of two main monophyletic groups in *Hypostomus*, with a division based on chromosome number. Robertsonian rearrangements were responsible for the formation of different diploid numbers and a diversity of karyotype formulas, as well as the cytogenetic marker patterns. Groups based on taxonomy were not rescued by gene analyses in this study, as they do not form monophyletic clades. NORs are predominantly multiple and located on st/a chromosomes, as well as in 18S rDNA sites. 5S rDNA sites are mostly observed in the interstitial position, following the trend already described for vertebrates.

## Acknowledgments

This work was supported by the Coordenação de Aperfeiçoamento de Pessoal de Nível Superior (CAPES – Fellowship for M.Sc. degree, www.capes.gov.br) and Conselho Nacional de Desenvolvimento Científico e Tecnológico (CNPq, 484626/2013-2, www.cnpq.br).

## Data Availability

The datasets generated and/or analysed during the current study are available from the corresponding author on reasonable request. Molecular sequence data will be deposited in GenBank after acceptance of the manuscript, as well as phylogenetic data in an appropriate database.

## Conflict of Interest

The authors declare that they have no conflict of interest.

## Ethical approval

All applicable international, national, and/or institutional guidelines for the care and use of animals were followed.

## Author contributions

Conceptualization: [Dinaíza Abadia Rocha-Reis], [Rubens Pasa], [Karine Frehner Kavalco]; Methodology: [Dinaíza Abadia Rocha-Reis]; Formal analysis and investigation: [Dinaíza Abadia Rocha-Reis], [Rubens Pasa], [Karine Frehner Kavalco]; Writing – original draft preparation: [Dinaíza Abadia Rocha-Reis]; Writing – review and editing: [Dinaíza Abadia Rocha-Reis], [Rubens Pasa], [Karine Frehner Kavalco]; Funding acquisition: [Karine Frehner Kavalco]; Supervision: [Rubens Pasa], [Karine Frehner Kavalco].

## Notes

### Competing Interest Statement

The authors have declared no competing interest.

