## Appendix B for "High congruence of karyotypic and molecular data on *Hypostomus* species from the Paraná River basin"

**Appendix B:** Details of the specimens collected and used in the study.

| Sample number | Laboratory record | Species | Collect point | Coordinates | | Hydrographic basin |
|---|---|---|---|---|---|---|
| 2 | 1702 | *H.* aff. *ancistroides* | São Miguel Arcanjo | 23°54’14.30”S | 47°55’46.75”O | Paranapanema river |
| 3 | 1705 | *H.* aff. *ancistroides* | São Miguel Arcanjo | 23°54’14.30”S | 47°55’46.75”O | Paranapanema river |
| 4 | 1726 | *H. ancistroides* | Araras | 22°22’59.64”S | 47°25’49.50”O | Tietê river |
| 5 | 1744 | *H.* aff. *ancistroides* | São Miguel Arcanjo | 23°54’44.58”S | 47°57’40.50”O | Paranapanema river |
| 6 | 1748 | *H. regani* | Terra Roxa | 20°43’34.00”S | 48°19’15.80”O | Grande river |
| 7 | 1749 | *H. regani* | Terra Roxa | 20°43’34.00”S | 48°19’15.80”O | Grande river |
| 8 | 1750 | *H. regani* | Terra Roxa | 20°43’34.00”S | 48°19’15.80”O | Grande river |
| 9 | 1751 | *H. margaritifer* | Terra Roxa | 20°43’34.00”S | 48°19’15.80”O | Grande river |
| 12 | 1762 | *H. regani* | Campina do Monte Alegre | 23°32’59.04”S | 48°30’44.58”O | Paranapanema river |
| 13 | 1801 | *H.* aff. *ancistroides* | Colina | 20°68’48.20”S | 48°53’14.20”O | Grande river |
| 14 | 1802 | *H.* aff. *ancistroides* | Colina | 20°68’48.20”S | 48°53’14.20”O | Grande river |
| 16 | 1901 | *H. ancistroides* | Araras | 22°22’59.64”S | 47°55’46.75”O | Tietê river |
| 17 | 1903 | *H. ancistroides* | Botucatu | 22°52’29.15”S | 48°22’27.50”O | Tietê river |
| 18 | 1907 | *H. strigaticeps* | Botucatu | 22°52’29.15”S | 48°22’27.50”O | Tietê river |
| 19 | 1908 | *H. ancistroides* | Botucatu | 22°52’29.15”S | 48°22’27.50”O | Tietê river |
| 20 | 1912 | *H. ancistroides* | Conchas | 23°00’03.00”S | 48°00’00.00”O | Tietê river |

| | | | | | | |
|---|---|---|---|---|---|---|
| 21 | 1914 | *H. ancistroides* | Conchas | 23°00’03.00”S | 48°00’00.00”O | Tietê river |
| 22 | 1918 | *H. ancistroides* | Indaiatuba | 23°05’39.12”S | 47°15’38.16”O | Tietê river |
| 23 | 1922 | *H. ancistroides* | Indaiatuba | 23°05’39.12”S | 47°15’38.16”O | Tietê river |
| 24 | 1923 | *H. ancistroides* | Indaiatuba | 23°05’39.12”S | 47°15’38.16”O | Tietê river |
| 25 | 1926 | *H. ancistroides* | Conchas | 23°00’03.00”S | 48°00’00.00”O | Tietê river |
| 26 | 1927 | *H. ancistroides* | Conchas | 23°00’03.00”S | 48°00’00.00”O | Tietê river |
| 27 | 1930 | *H. ancistroides* | Terra Roxa | 20°43’34.00”S | 48°19’15.80”O | Grande river |
| 28 | 1932 | *H. ancistroides* | Terra Roxa | 20°43’34.00”S | 48°19’15.80”O | Grande river |
| 29 | 2440 | *Hypostomus sp.* | Uberabinha river | 18°40’40.37”S | 48°30’23.44”O | Paranaíba river |
| 30 | 2441 | *Hypostomus sp.* | Uberabinha river | 18°40’40.37”S | 48°30’23.44”O | Paranaíba river |
| 31 | 2449 | *H. regani* | Paranaíba river P2 | 19°04’32.08”S | 46°24’33.07”O | Paranaíba river |
| 32 | 2471 | *H. regani* | Paranaíba river P3 | 18°40’00.51”S | 46°31’13.12”O | Paranaíba river |
| 33 | 2548 | *H.* cf. *tietensis* | Uberabinha river | 18°40’40.37”S | 48°30’23.44”O | Paranaíba river |
| 34 | 2585 | *H. strigaticeps* | Meia Ponte river | 17°44’13.15”S | 49°25’52.24”O | Paranaíba river |
| 35 | 2587 | *H. faveolus* | Meia Ponte river | 17°44’13.15”S | 49°25’52.24”O | Paranaíba river |
| 36 | 2588 | *H. paulinus* | Meia Ponte river | 17°44’13.15”S | 49°25’52.24”O | Paranaíba river |
| 37 | 2589 | *H. strigaticeps* | Meia Ponte river | 17°44’13.15”S | 49°25’52.24”O | Paranaíba river |
| 38 | 2590 | *Hypostomus* sp. 2 | Meia Ponte river | 17°44’13.15”S | 49°25’52.24”O | Paranaíba river |
| 39 | 2592 | *Hypostomus* sp. | Meia Ponte river | 17°44’13.15”S | 49°25’52.24”O | Paranaíba river |
| 40 | 2593 | *H. ancistroides* | Meia Ponte river | 17°44’13.15”S | 49°25’52.24”O | Paranaíba river |

| | | | | | | |
|---|---|---|---|---|---|---|
| 41 | 2646 | *H. ancistroides* | Uberabinha river | 18°40’40.37”S | 48°30’23.44”O | Paranaíba river |
| 42 | 2680 | *Hypostomus* sp. | Uberabinha river | 18°40’40.37”S | 48°30’23.44”O | Paranaíba river |
| 43 | 2681 | *Hypostomus* sp. | Paranaíba river P1 | 19°10’56.23”S | 46°19’58.06”O | Paranaíba river |
| 44 | 2686 | *Hypostomus* sp. | Uberabinha river | 18°40’40.37”S | 48°30’23.44”O | Paranaíba river |
| 45 | 2687 | *H. ancistroides* | Paranaíba river P1 | 19°10’56.23”S | 46°19’58.06”O | Paranaíba river |
| 46 | 2747 | *H. regani* | Paranaíba river P2 | 19°04’32.08”S | 46°24’33.07”O | Paranaíba river |
| 47 | 2748 | *H. regani* | Paranaíba river P2 | 19°04’32.08”S | 46°24’33.07”O | Paranaíba river |
| 48 | 2750 | *Hypostomus* sp. | Paranaíba river P2 | 19°04’32.08”S | 46°24’33.07”O | Paranaíba river |
| 49 | 2751 | *Hypostomus* sp. | Paranaíba river P2 | 19°04’32.08”S | 46°24’33.07”O | Paranaíba river |
| 50 | 2776 | *Hypostomus* sp. | Paranaíba river P3 | 18°40’00.51”S | 46°31’13.12”O | Paranaíba river |
| 51 | 2778 | *H. regani* | Paranaíba river P3 | 18°40’00.51”S | 46°31’13.12”O | Paranaíba river |
| 52 | 2779 | *Hypostomus* sp. | Paranaíba river P3 | 18°40’00.51”S | 46°31’13.12”O | Paranaíba river |
| 53 | 2800 | *Hypostomus* sp. | Água Grande | 19°13’49.31”S | 46°12’20.70”O | Paranaíba river |
| 54 | 2801 | *Hypostomus* sp. | Água Grande | 19°13’49.30’”S | 46°12’20.70”O | Paranaíba river |
| 55 | 2880 | *Hypostomus* sp. | São João river | 19°18’29.72”S | 46°23’49.62”O | Paranaíba river |
| 56 | 2888 | *Hypostomus* sp. | São João river | 19°18’29.72”S | 46°23’49.62”O | Paranaíba river |
| 57 | 2983 | *Hypostomus* sp. | São João river | 19°18’29.72”S | 46°23’49.62”O | Paranaíba river |
| 58 | 3064 | *Hypostomus* sp. | Paranaíba river P4 | 19°07’52.57”S | 46°23’30.42”O | Paranaíba river |
| 59 | 3065 | *H. ancistroides* | Paranaíba river P4 | 19°07’52.57”S | 46°23’30.42”O | Paranaíba river |
| 61 | 3103 | *H. ancistroides* | Paranaíba river P1 | 19°10’56.23”S | 46°19’58.06”O | Paranaíba river |

| | | | | | | |
|---|---|---|---|---|---|---|
| 62 | 3112 | *H. regani* | Paranaíba river P3 | 18°40’00.51”S | 46°31’13.12”O | Paranaíba river |
| 63 | 3117 | *Hypostomus* sp. | Paranaíba river P5 | 18°55’04.26”S | 46°30’20.68”O | Paranaíba river |
| 64 | 3123 | *H. strigaticeps* | Paranaíba river P5 | 18°55’04.26”S | 46°30’20.68”O | Paranaíba river |
| 65 | 3125 | *H. regani* | Paranaíba river P5 | 18°55’04.26”S | 46°30’20.68”O | Paranaíba river |
| 66 | 3128 | *H. regani* | Paranaíba river P5 | 18°55’04.26”S | 46°30’20.68”O | Paranaíba river |
| 67 | 3140 | *H. regani* | Paranaíba river P5 | 18°55’04.26”S | 46°30’20.68”O | Paranaíba river |
| 68 | 3184 | *Hypostomus* sp. | Paranaíba river P5 | 18°55’04.26”S | 46°30’20.68”O | Paranaíba river |
| 69 | 3205 | *H. regani* | Paranaíba river P1 | 19°10’56.23”S | 46°19’58.06”O | Paranaíba river |
| 70 | 3206 | *H. regani* | Paranaíba river P3 | 18°40’00.51”S | 46°31’13.12”O | Paranaíba river |
