## Appendix C for "High congruence of karyotypic and molecular data on *Hypostomus* species from the Paraná River basin"

**Appendix C:** Sequences of *Hypostomus* and *Pterygoplichthys* (outgroup) species used in the analyzes. Available on GenBank.

| Accession | Species | Gene | Accession | Species | Gene |
|---|---|---|---|---|---|
| KC170033 | *Pterygoplichthys disjunctivus* | *mt-co1* | HM404959 | *Pterygoplichthys etentaculatus* | *mt-co1* |
| KP959938 | *Pterygoplichthys gibbiceps* | *rag1* | HQ267783 | *Pterygoplichthys joselimaianus* | *mt-cyb* |
| GQ225501 | *Pterygoplichthys pardalis* | *rag2* | DQ133773 | *Pterygoplichthys zuliaensis* | *mt-cyb* |
| HM376401<br>GQ225397 | *Hypostomus cochliodon* | *mt-co1* | GU702243<br>GU702254 | *Hypostomus affinis* | *mt-co1* |
| GU701719<br>JN988938 | *Hypostomus commersoni* | *mt-co1* | GU701713<br>JN988940 | *Hypostomus derbyi* | *mt-co1* |
| KJ720692<br>JN026849 | *Hypostomus plecostomus* | *mt-co1* | JN988952<br>GU701659 | *Hypostomus topavae* | *mt-co1* |
| GU701668<br>GU701667 | *Hypostomus strigaticeps* | *mt-co1* | GU701681<br>GU701683 | *Hypostomus paulinus* | *mt-co1* |
| GU701703<br>GU701707 | *Hypostomus hermanni* | *mt-co1* | GU701733<br>GU701708 | *Hypostomus heraldoi* | *mt-co1* |
| GU701728<br>GU701731 | *Hypostomus albopunctatus* | *mt-co1* | GU701699<br>JN988944 | *Hypostomus iheringii* | *mt-co1* |
| GU701686<br>GU701689 | *Hypostomus nigromaculatus* | *mt-co1* | | | |
